# FunCLAN: A novel and generalisable method for categorising protein complex conformations, measuring their similarities, and solving the chain equivalence problem

**DOI:** 10.1101/2025.06.16.659920

**Authors:** Daniel Celis Garza, Joseph I. J. Ellaway, Sri Devan Appasamy, Eugene Krissinel, Sameer Velankar

## Abstract

We present FunCLAN, a novel method for finding, categorising, and measuring conformational relationships between protein complexes, as well as for finding corresponding chain pairs between them. Its purpose is to provide a general tool, applicable to a broad range of complexes. As such, it only utilises sequence similarity and superposition transformations to compare rigid bodies (chains), and how they transform with relation to each other. This lets us clusterise transformations, which lets us identify sets of similar and dissimilar transformations across the dataset. We use this information to classify conformations according to how the transformations relate to each other at chain and complex level. We have also included two data, and scale agnostic methods that can make informed predictions as to the number of clusters (conformers) present in the dataset. We also present a way to superpose entire complexes using the alignment and chain correspondence data for visualisation purposes. The entire process can be generalised to domains and even single chains, but this requires bespoke/specialised preprocessing of the data as well as prior knowledge of the structures to be analysed. This is out of the scope of this paper, but we outline a method to do so in the supplementary information.

## Main

Large macromolecular machines are flexible molecules that change shape or conformations to function. Understanding their dynamic nature is vital for gaining mechanistic insights into the function and evolution of these macromolecular machines and their role in complex biological processes. The dynamic nature of individual components of these macromolecular machines is equally important in forming the complexes and, in many cases, for the regulation of these molecules, such as switching between active and inactive forms for enzymatic activity [15]. Molecular structures and the details of the different conformational states at atomic resolution can also help explain the effects of genetic variants [14] facilitate the drug design process, elucidate the mechanism of drug resistance and help in protein design [27].

Apart from the traditional molecular dynamics simulation, rapid advances in experimental and structure prediction methods are providing ways for studying the dynamic behaviour of macromolecules. The availability of advanced electron microscopy, time resolved and X-ray free electron laser (XFEL) methods can provide atomic details of the structure dynamics. The recent advances in AI methods to analyse cryo-electron tomography (cryo-ET) data is uncovering dynamics of large macromolecular complexes in their natural environment inside cells [13.

Established in 1971, the Protein Data Bank (PDB) [21] is a single global archive of macromolecular structure. With over 220,00 experimentally determined structures of macromolecules and their complexes with ligands, other proteins or nucleic acid, the PDB provides a rich source of information for the study of protein structure and function. The PDB contains many experimentally determined macromolecular complex structures in multiple forms, typically bound to different ligands, crystallised in different space groups or studied using electron microscopy under different sample conditions, in complex with different cofactors, different sequence variants, different oligomeric states, and others providing an array of individual snapshot of conformations. The multiple conformational states for individual macromolecules present in the PDB can be exploited to characterise the high-dimensional protein conformational landscapes. Characterisation of the different conformational states can lead to better understanding of the functional mechanism by revealing conserved domains and principal conformation points, like hinges, providing a way to uncover underlying principles of conformational patterns.

Such an analysis can make use of a robust software tool for aligning the different macromolecular structures to reveal the similarities and differences between the different conformational states. In the impressive variety of software tools developed for structural alignment of protein chains, only a few (e.g. ProSMART [12], GESAMT [5], FATCAT [23] do not consider protein chains as rigid bodies and, therefore, can be used for inferring conformational states from polypeptide collections. To our best knowledge, other than Vast+ [9], which has a different use case than *FunCLAN*, there are no tools that interrogate the conformational landscape of complexes in a similar manner. However, complexes represent particular biological interest as the physiological activity of many (if not most) proteins is associated with their oligomeric form. Technically, the analysis of whole complex conformations is complicated by the fact that they cannot be validly presented as polypeptide “super-chains”. Chains must be treated independently or as units to be analysed. Therefore, straightforward application of existing methods based on monomeric proteins is not feasible.

We present FunCLAN (Functional annotations through Conformational Landscape Analysis), a new algorithm for the identification of clusters of chains, and complexes with similar conformational characteristics, superposition of protein complexes, and identification of corresponding chain pairs between conformers. FunCLAN is designed to quantify the similarities and differences in how macromolecular structures transform between conformations, allowing it to cluster and classify the structures to identify different conformational states via unsupervised machine learning. Although FunCLAN was developed specifically for studying concerted movements of large complexes, the approach can be generalised to classify single chains and protein-ligand binding sites. Given a set of n>1 structurally similar complexes, FunCLAN finds correlations in relative orientation of their chains, then scores and clusters them.

If an experimentally obtained dataset is viewed as a series of snapshots of a dynamic complex, FunCLAN analysis shows which parts of the complex move coherently and to which degree the coherency is maintained. FunCLAN does not make assumptions on the relative spatial position of the moving parts being compared and can detect relative movement between individual components that may not have direct interaction. The results are represented as distance matrices summarised by dendrograms. Other outputs include the compressed score matrices, chain correspondences between complex samples, aligned sequences, chain-chain superpositions, and whole complex superpositions. The algorithm is implemented as an open-source C++/Python tool under the Apache 2.0 licence and is freely available from https://gitlab.com/dcelisgarza/FunCLAN/-/tree/main.

## Results

### Algorithm Overview

*FunCLAN* is an algorithm that exploits the observation that while superposition matrices provided by the conventional methods between equivalent chains of non-conformed, “identical” (up to some tolerance), complexes are all equal to each other, the relationships between sets of transformations are not necessarily identical. For example, suppose that chains *A*, *B* of complex Θ_1_ correspond to chains *C*, *D* of complex Θ_2_, and their Pairwise Superposition Matrices (PSMs) are *T*(*A*|*C*) and *T*(*B*|*D*), respectively. Then the scale of similarity between Θ_1_ and Θ_2_ can be measured by the PSM difference Δ*T*(*A*|*B*, *C*|*D*) = |*T*(*A*|*C*) − *T*(*B*|*D*)|. In general, the conformational landscape of multimeric complexes of size *N* > 2 may be described by groups of chains forming conserved sub-structures, with a degree of variability corresponding to the range of Δ*T* values within the groups.

FunCLAN starts from calculating PSMs between all similar chains in a given set of homologous complexes. The similarity is identified by a sequence identity threshold, which is acceptable as FunCLAN is designed to operate on highly homologous structures. If a complex includes several chains with a given sequence, several PSMs correspond to each of them, which can be clustered in *n*_*sym*_ ≤ *N* well-packed clusters of size *N*, where *n*_*sym*_ is the complex’s symmetry number. Each such cluster corresponds to an alternative (and equivalent) overall complex superposition. To resolve this ambiguity, FunCLAN leaves only PSMs found in the best-packed cluster in consideration. FunCLAN identifies coherently conforming groups of chains by hierarchical clustering of PSM differences Δ*T*, which results in a dendrogram representing hierarchical association of chains in the complex. It then performs a further clusterisation using a distances of distances approach proposed in [7], which has a number of desirable characteristics, though both whole-complex clusterings can be useful.

The concept of FunCLAN analysis is illustrated in Figure 1 using a schematic complex that forms three conformers (Fig. 1a). The dendrogram for conformers 1 and 3 (Fig. 1b) identifies two sets of chains with conserved relative positions between these conformers. Similarly, the dendrogram for conformers 2 and 3 (Fig. 1c) indicates that the circle, triangle, and hexagonal chains largely maintain their spatial relationships, whereas the pentagonal chains exhibit the most variation.

**Figure 1.**
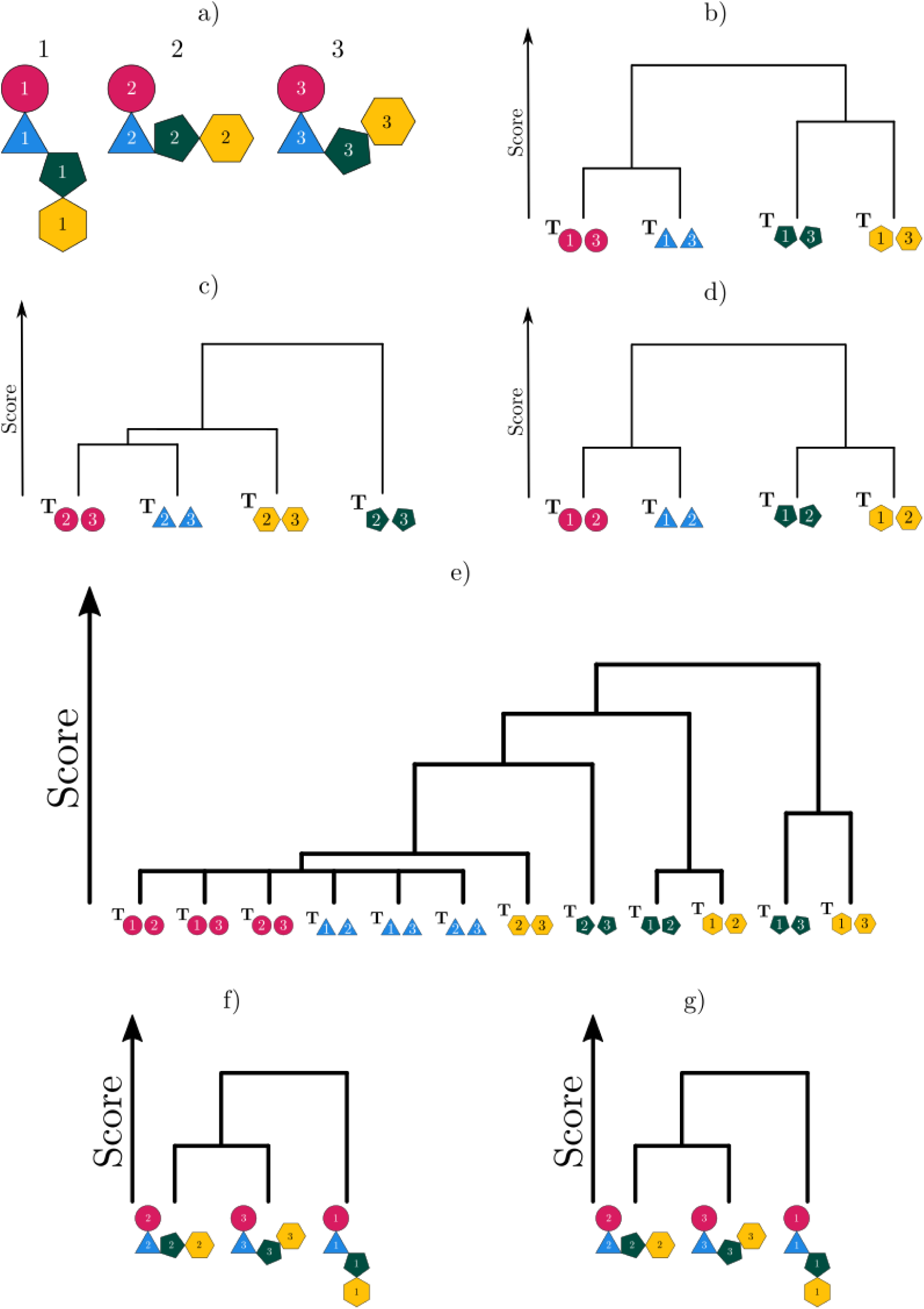
**a)** Schematic representation of a complex consisting of four chains, distinguished by different shapes, colours, and labels. Three conformers are shown to illustrate the analysis discussed in the main text. **b)** FunCLAN dendrogram comparing conformers 1 and 3 from (a). **c)** FunCLAN dendrogram comparing conformers 2 and 3 from (a). **d)** FunCLAN dendrogram comparing conformers 1 and 2 from (a). **e)** FunCLAN chain-level dendrogram incorporating all conformers from (a), showing hierarchical chain associations. **f)** FunCLAN whole-complex dendrogram for all three conformers in (a), summarizing the distance matrix underlying (e). **g)** Dendrogram obtained after clustering second-order distances (distances of distances). Since there are only three conformers in this example, its structure closely resembles (e), though the scores differ.

The dendrogram for conformers 1 and 2 (Fig. 1d) reveals two distinct sub-complexes: (1) circles and triangles, and (2) pentagons and hexagons. While these sub-complexes are preserved across conformers, their mutual orientations vary. Fig. 1e can be seen as a projection of the overall conformation landscape of all three conformers. This hierarchical clustering of chain associations enables the identification of potential sub-complexes based on a chosen score threshold. Higher scores indicate weaker associations, reflecting greater variability in relative positions within sub-complexes.

Additionally, the dendrogram of chain associations can be transformed into a dendrogram that classifies the conformers as a whole (Fig. 1f). This representation reveals the degree of similarity between conformers through pairwise comparisons. The classification can be further refined by comparing each row of the distance matrix to all other rows used to construct Fig. 1f, providing a conformer-centric view of the complex’s conformational space. In this particular case, the dendrograms in Figs. 1f and 1g are nearly identical, with differences limited to score variations due to the small number of conformers. However, as demonstrated in subsequent sections, these differences can be significant in biologically meaningful changes in assembly structure.

### Example 1. SARS-CoV2 Spike Protein

One of the most clinically significant and widely studied proteins since 2020 is the SARS-CoV-2 spike protein, which played a central role in the COVID-19 pandemic. This trimeric glycoprotein extends from the viral surface and is essential for viral entry into host cells. By binding to the human angiotensin-converting enzyme 2 (ACE2) receptor, the spike protein initiates the fusion process, enabling the virus to penetrate host cells and hijack their machinery for replication [2,16,17,19,24].

A key characteristic of the SARS-CoV-2 spike protein is its structural plasticity, which directly impacts its function. The spike protein transitions between multiple conformations essential for viral fusion and immune evasion. Each protomer contains a receptor-binding domain (RBD), which can adopt one of two key states: an “up” (open) conformation, allowing ACE2 receptor binding, or a “down” (closed) conformation, concealing the RBD from neutralizing antibodies. Typically, one or two RBDs adopt the open state at any given time, while the rest remain closed, balancing receptor engagement with immune evasion.

However, subtle structural changes – less apparent than the movement of individual RBD domains – can have a greater impact on the interactions between spike protein subunits. This raises an intriguing question: how do we define a conformer?

### Direct comparison of conformations

Fig. 2a illustrates the relationships between all matched sets of chains in the dataset, revealing a clear underlying structure in their interactions. It is the primary data that FunCLAN uses to derive its results. Figs 2b and 2d (1-5) reveal five distinct conformational groups, summarized in Table 1. As may be seen in the table, while the notations used by different authors (Variant, Conformation, and State) are not fully consistent across the dataset, the clustering results generally align with the existing classifications where applicable. Notably, when analyzing the superposition of the entire complex, distinct structural commonalities emerge within each cluster.

**Figure 2.**
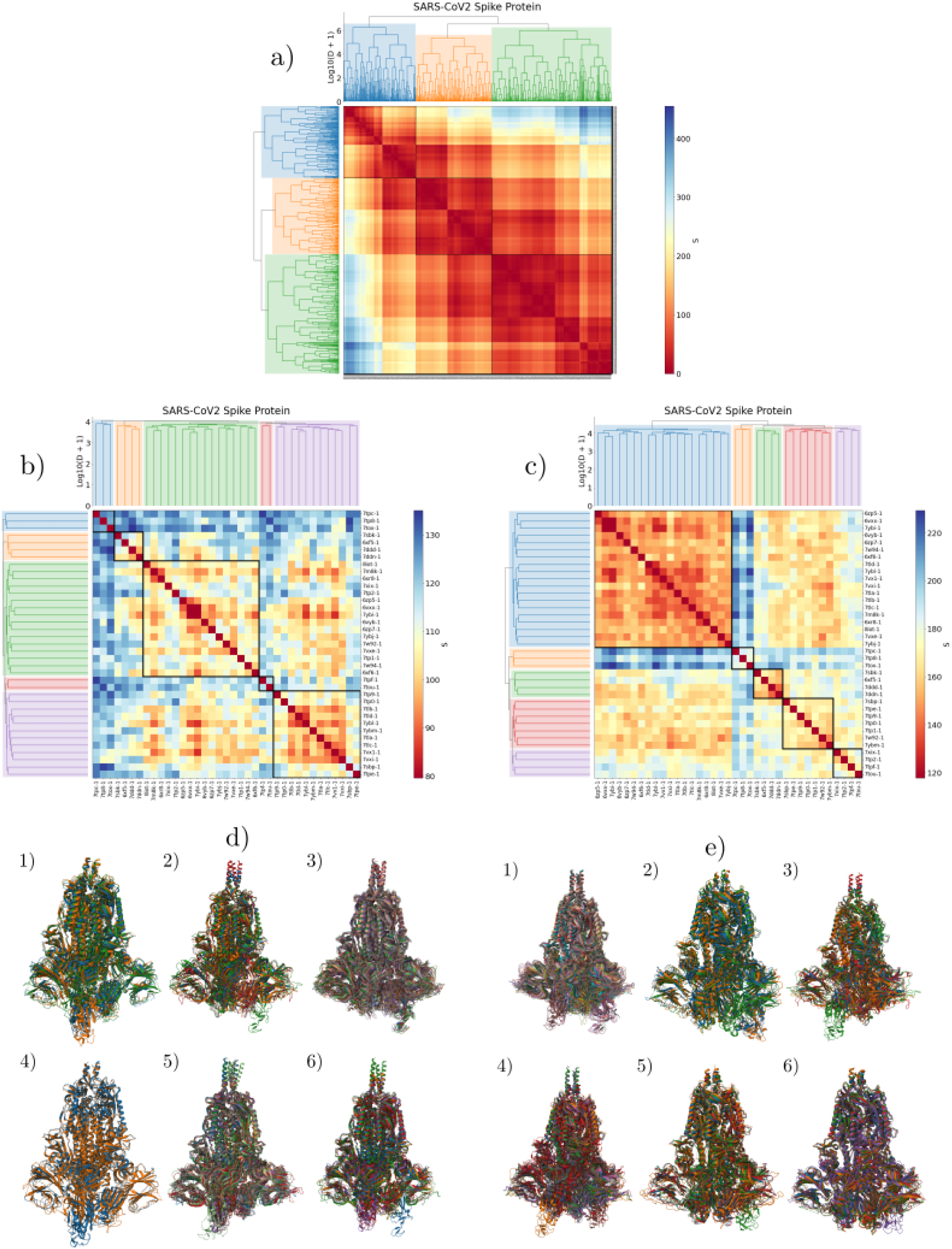
**a)** Chain level conformational landscape. Number of clusters determined via the second-order [24] difference statistic. **b)** SARS-CoV2 spike protein score matrix and dendrogram. Number of clusters determined via the second-order difference statistic, clusters are taken to be 1-5 from top left to bottom right. **c)** SARS-CoV2 spike protein distances of distances matrix and dendrogram. Number of clusters determined via the standardised silhouette score [8], clusters are taken to be 1-5 from top left to bottom right. **d)** From top left to bottom right they are the superposed structures from clusters **d1)**, **d2)**, **d3)**, **d4)** and **d5)**. **d6)** is a superposition of one representative from each cluster. **e)** From top left to bottom right they are the superposed structures from clusters **e1)**, **e2)**, **d3)**, **e4)** and **e5)**. **e6)** is a superposition of one representative from each cluster. A cluster’s representative is the structure whose score is closest to the median score of the cluster. All structures were taken to a common frame with the same viewing angle. The last superposition is of a representative of each cluster. The colour scale of the matrices has been adjusted to exclude outliers more than a factor of 10 medians away from the median score of the entire matrix, because it makes it easier to visualise small score differences in score.

**Table 1.**
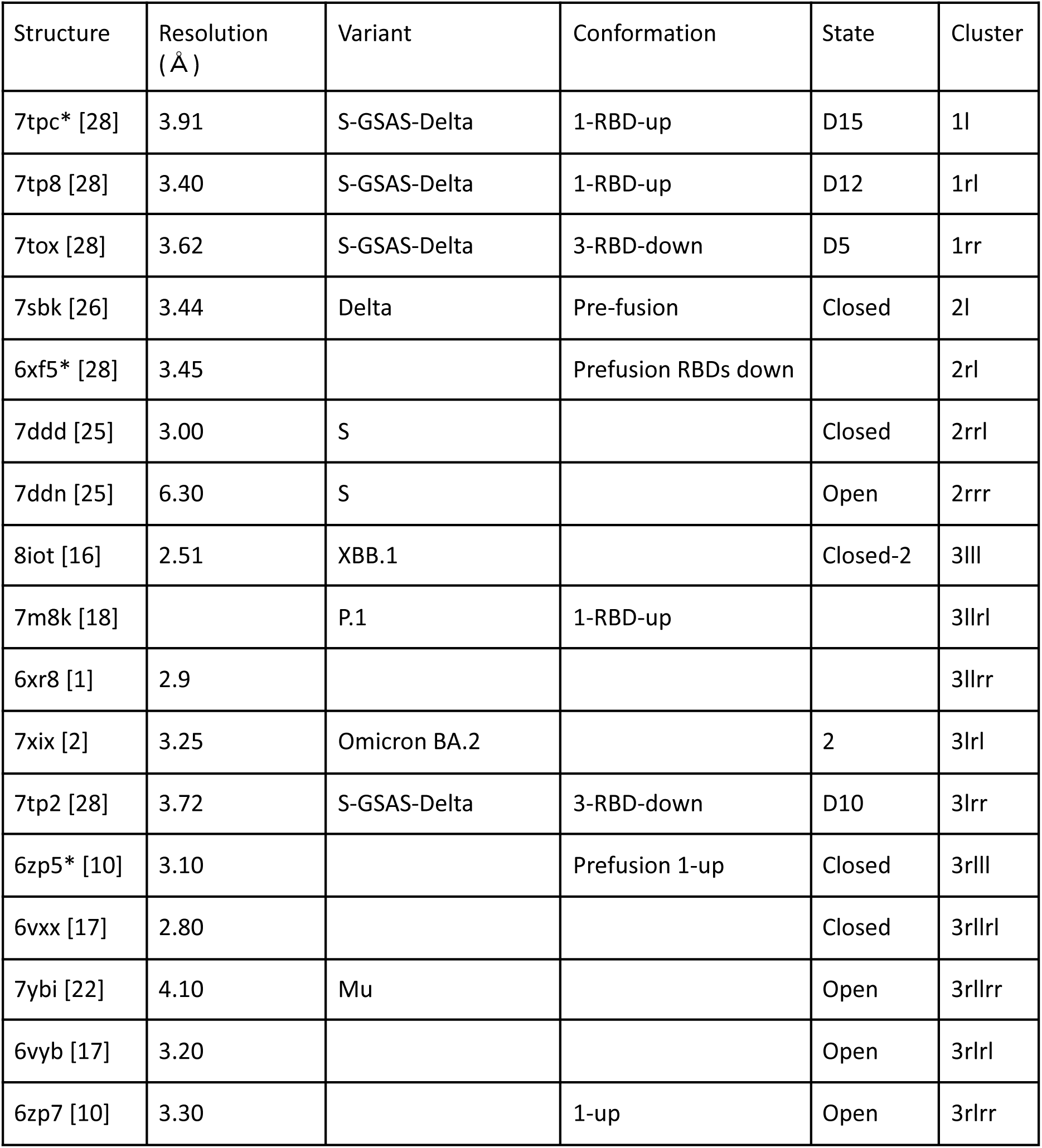
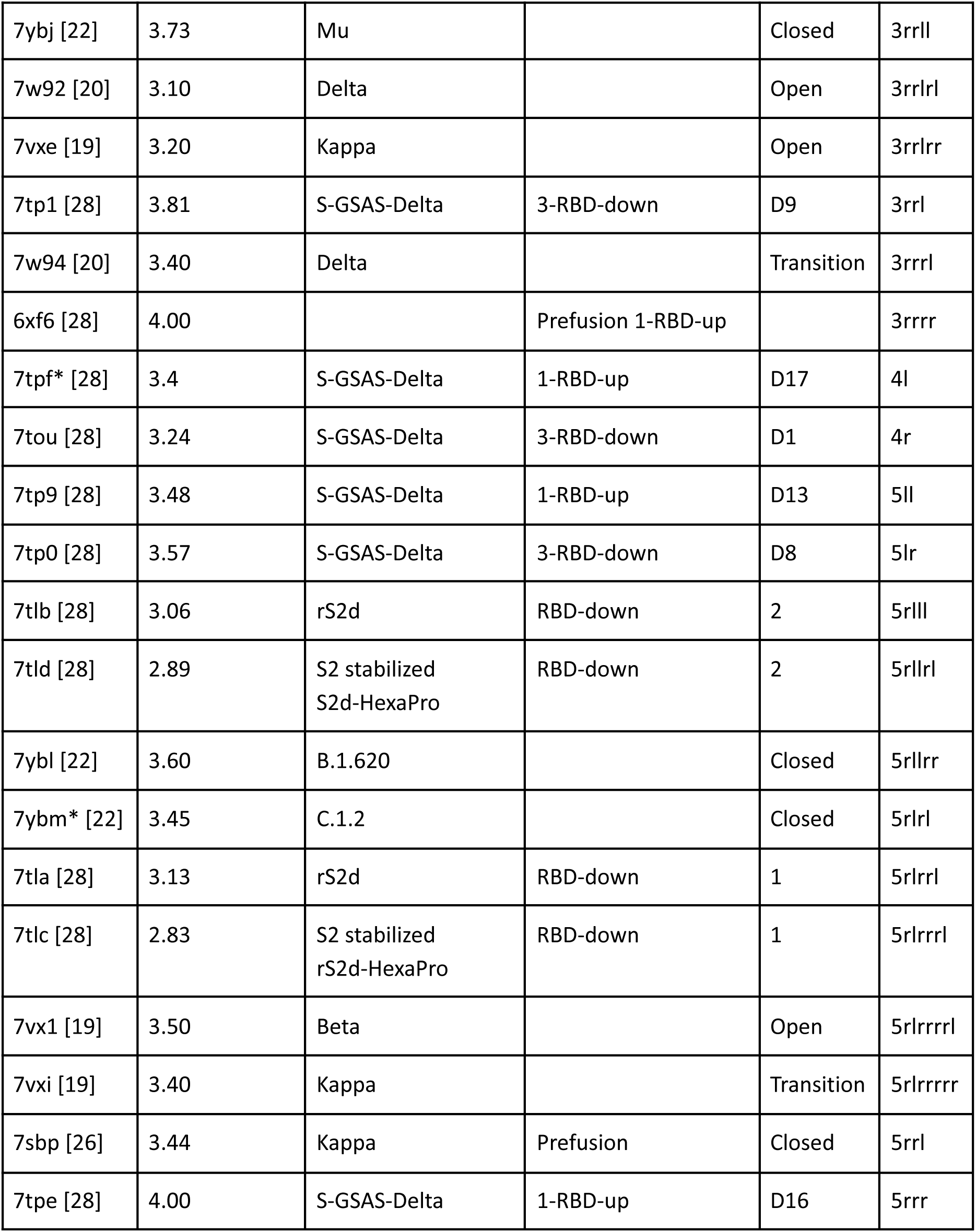
The table entries correspond to those in the dendrogram and distance matrix in Figs. 2b, 2d. Square brackets indicate the source publication of each structure. Cluster keys denote hierarchical clustering from left to right. The “Variant,” “Conformation,” and “State” columns are taken from PDB entries. The “Cluster” row shows each structure’s position in the dendrogram, with numbers representing clusters (left to right) and “l” or “r” indicating branch direction. For example, 7tpe belongs to the fifth cluster and follows right-side branches to its leaf. Leaf positions may swap without altering dendrogram meaning. Asterisks (*) mark characteristic cluster representatives, selected via a doubly-centered median score (see Methods).

It is important to recognize that clustering algorithms provide an approximation of relationships between objects rather than a definitive classification. The full relationship is more accurately represented in the distance matrix (Fig, 2a-c). Additionally, the superposition images in Figs. 2d and 2e are captured from a single angle, with many structural features obscured due to the number of overlapping structures. As a result, visual assessments of similarity or dissimilarity from superpositional images should not be taken as absolute. Nevertheless, broad characteristic patterns can still be discerned within each cluster.

Thus, analyzing Figs. 2b and 2d, we observe that clusters 1 and 4 consist of samples that are similarly distant from all other samples. In fact, one could argue that they might form a distinct cluster of their own. This similarity is further supported by the data in Table 1. We also observe that the complexes in cluster 2 are largely similar to one another, aside from a flexible domain in one of the structures. This exemplifies a key feature of FunCLAN: rather than focusing on changes within individual chains, it prioritizes the relationships between them. Next, the complexes in cluster 3 appear similar to those in cluster 2, though they are slightly more compact. However, this is a case where perspective obscures key differences, so we encourage readers to explore the relevant *.msvj files provided in the supplementary information, which can be interactively viewed in a web browser. Notably, one complex in cluster 3 contains a chain with a flexible domain strikingly similar to the outlier from cluster 2. This further illustrates the logic behind FunCLAN and serves as motivation for a deeper analysis using distances of distances, examined in the following subsection. Lastly, the structures in cluster 5 exhibit very low intra-cluster variability, indicating a high degree of internal consistency. While distinct from the other clusters, the relationships within this group remain well-defined.

Overall, we conclude that the clusters largely align with the author annotations in Table 1. The distance matrix in Fig. 2b demonstrates that clusters with high internal similarity exhibit greater consistency in their superpositions in Fig. 2d. When examining the superposed structures in Fig. 2d, each cluster displays distinct features, with differences between clusters being visually apparent—except for clusters 1 and 4, which are more similar to each other. Given that the chosen viewpoint may obscure some structural details, we have provided *.msvj files in the supplementary information for interactive exploration.

## Distances of Distances

The previous results were obtained by directly comparing each sample to all others using their conformational coordinates. However, by applying the distances of distances approach from [7], we can analyze the entire phase space more comprehensively. To determine the optimal number of clusters, we used standardized silhouette scores [8] (Fig. 2c), as the second-order [24] difference statistic yielded only two clusters.

From the comparison of Figs. 2b and 2c, we can immediately observe that Fig. 2c contains fewer and less pronounced out-of-cluster hotspots. This is a desirable outcome, as it indicates that the clusters are more homogeneous and centered around a representative measure. Additionally, Fig. 2e presents a different perspective from Fig. 2d. This is intentional, as the clusters shown in superpositions (d1, e2) and (d2, e3) are the same, and providing an alternate viewpoint helps reveal different structural details. Since many clusters share similarities, viewing them from multiple angles enhances our ability to discern subtle differences. Table 2 provides a summary of Figs. 2c and 2e.

**Table 2:**
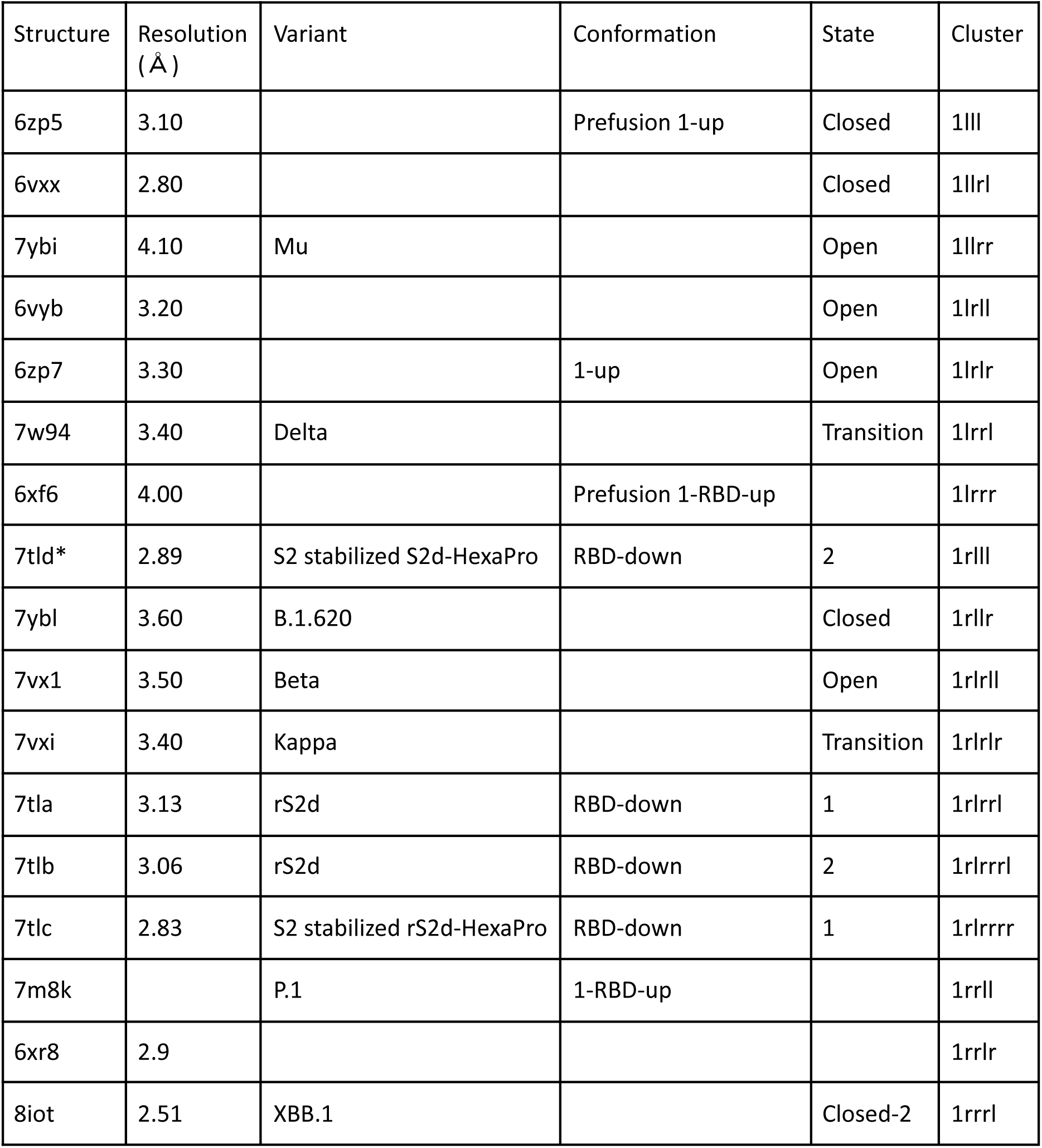
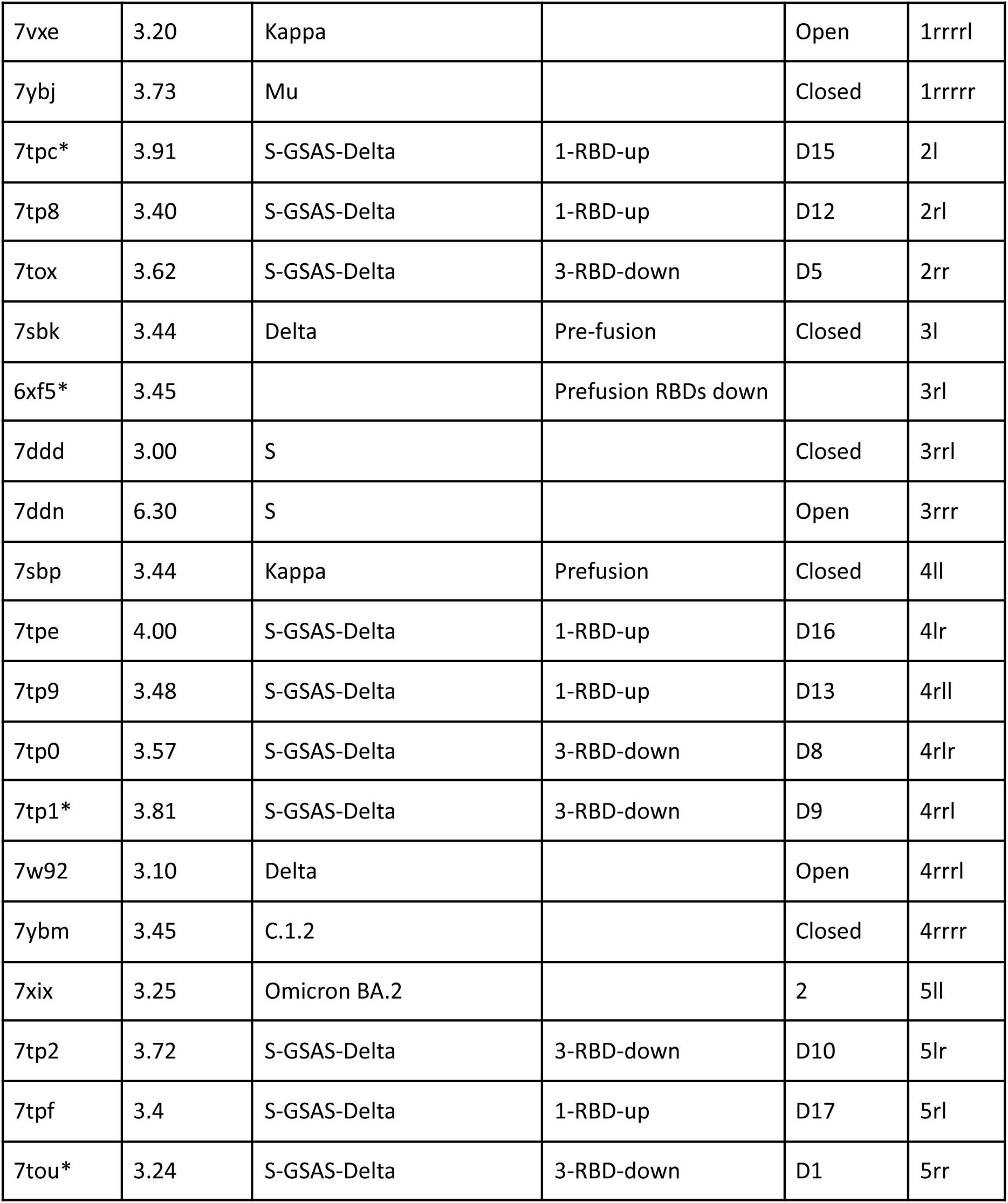
The table entries correspond to the entries in the dendrogram and distance matrix in Figs. 2c and 2e. The cluster keys are created from the cluster (from left to right) to which an entry belongs. All annotations follow the same methodology as in Table 1.

**Table 3.**
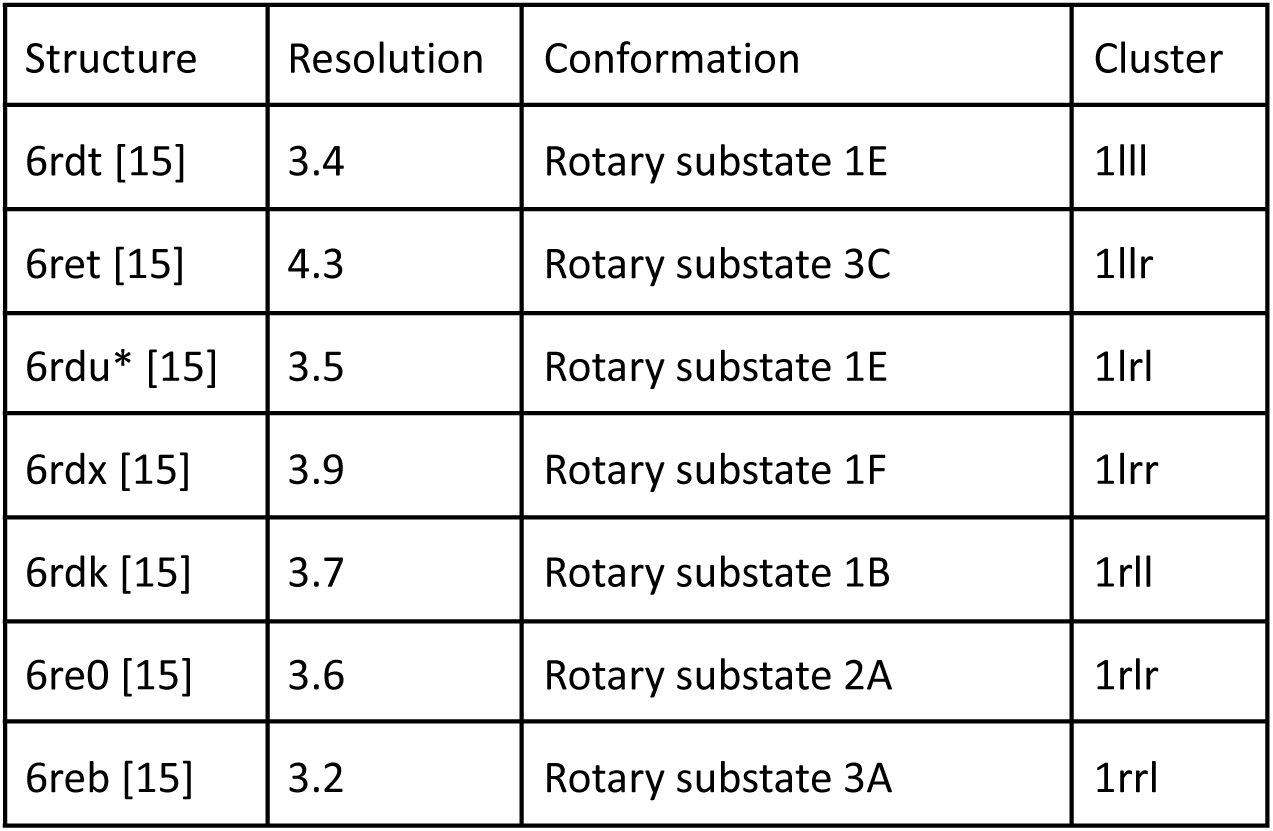
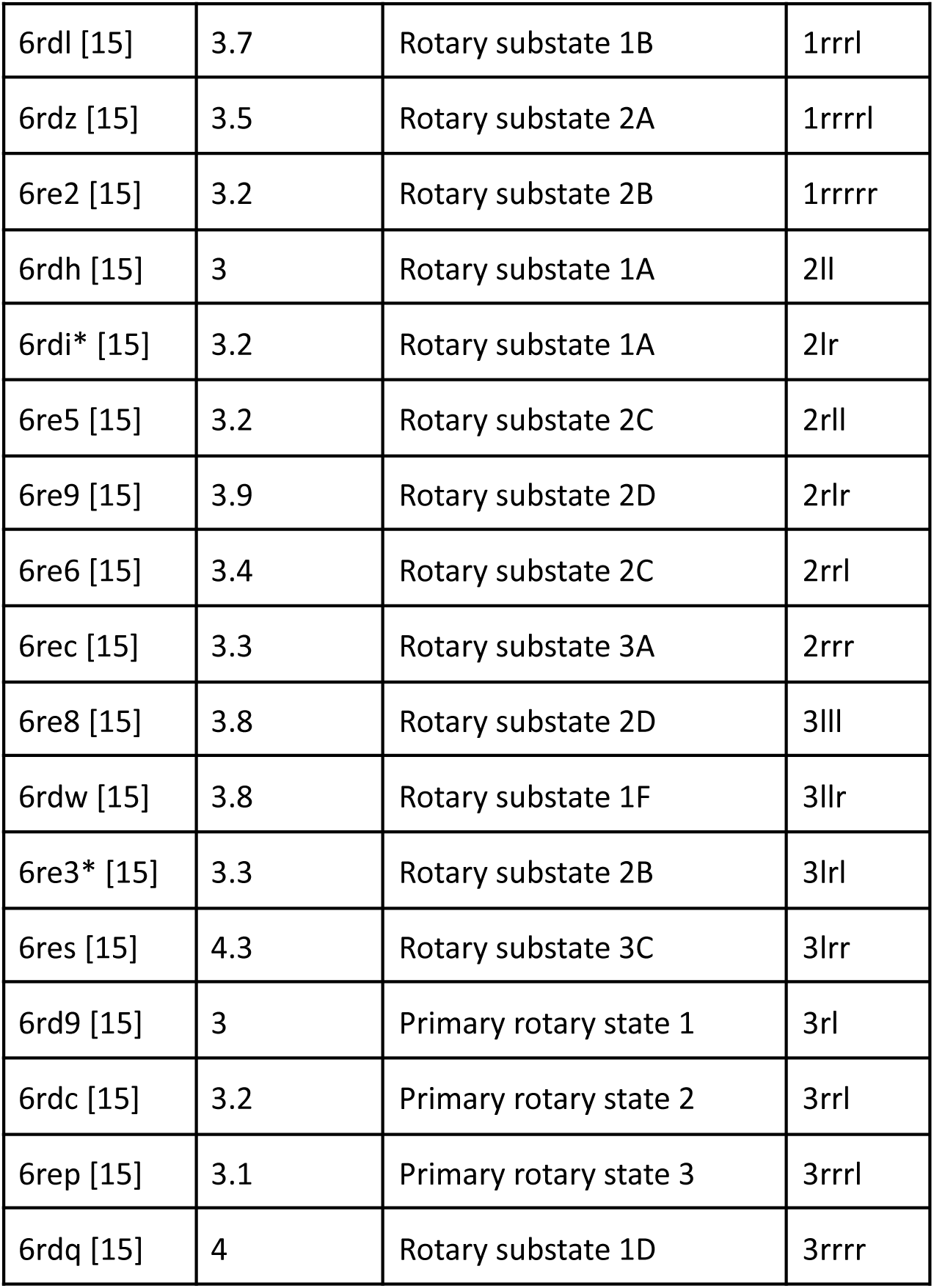
The table entries correspond to the entries in the dendrogram and distance matrix Fig. 3b and 3d. The cluster keys are created from the cluster (from left to right) to which an entry belongs. The “Resolution”, “Method” and “Conformation” headings come from each structure’s PDB entry. The cluster labels and representatives are labelled with the methods described in Table 1.

Overall, the clustering results from the “distances of distances” approach closely resemble the previous ones. Notably, clusters 1 and 2 from Fig. 2b corresponds to clusters 2 and 3 in Fig. 2c. However, some shifts are observed:

● 7xix and 7tp2 move from cluster 3 to cluster 5.
● 7w92 and 7tp1 move from cluster 3 to cluster 4.
● 7tpf and 7tou move from cluster 4 to cluster 5.
● 7tld, 7ybm, 7vxi, 7tla, 7tlb, and 7tlc move from cluster 5 to cluster 1.

Most of these changes involve structure pairs that were already classified as relatively similar, leading them to shift together.

In general, each cluster retains distinct characteristics, and one could argue that the “distances of distances” clustering method produces slightly more compact clusters. However, this is difficult to fully appreciate in a 2D image, making it more effective to explore these clusters interactively using the *.msvj files provided in the supplementary information.

### Example 2. Polytomella F-ATP synthase

ATP synthase is a vital enzyme located in the membranes of mitochondria, chloroplasts, and bacteria, where it plays a key role in cellular energy production. It generates adenosine triphosphate (ATP), the cell’s primary energy currency, by utilizing the energy from a proton gradient across the membrane. In Polytomella, F-ATP synthase forms a large heteromeric complex composed of 30 chains representing 18 distinct types. Within this structure, a homodecameric domain and two homotrimeric domains can be identified, highlighting its intricate organization and functional specialization.

Although this protein is traditionally described as having three rotary conformers, the reality appears far more complex. As seen in the *.msvj superposition files – and to some extent in the superposition images – there seems to be a continuum of conformations, particularly in how the side chains fan out. This underscores the same challenges observed in the SARS-CoV example but to an even greater degree.

#### Direct comparison of conformations

Figure 3a presents the relationships between all matched sets of chains in the dataset of 24 ATP synthase complexes used in this study. The dataset exhibits a structured organization in how these chains relate to one another. While ATP synthase is traditionally described as having three well-defined conformers, the clustering results reveal a more nuanced picture (cf. Figs. 3b and 3d). Although conformers generally align with their FunCLAN categorization, the distribution is not strictly uniform. For example, Cluster 1 primarily consists of rotary state 1 but also includes four instances of rotary state 2 alongside five of rotary state 1. Similarly, Cluster 2 is predominantly composed of rotary state 2, yet it contains two instances of rotary state 1 compared to three of rotary state 2. Cluster 3 also exhibits a mixture of rotary states, further highlighting the overlap between these categories.

**Figure 3.**
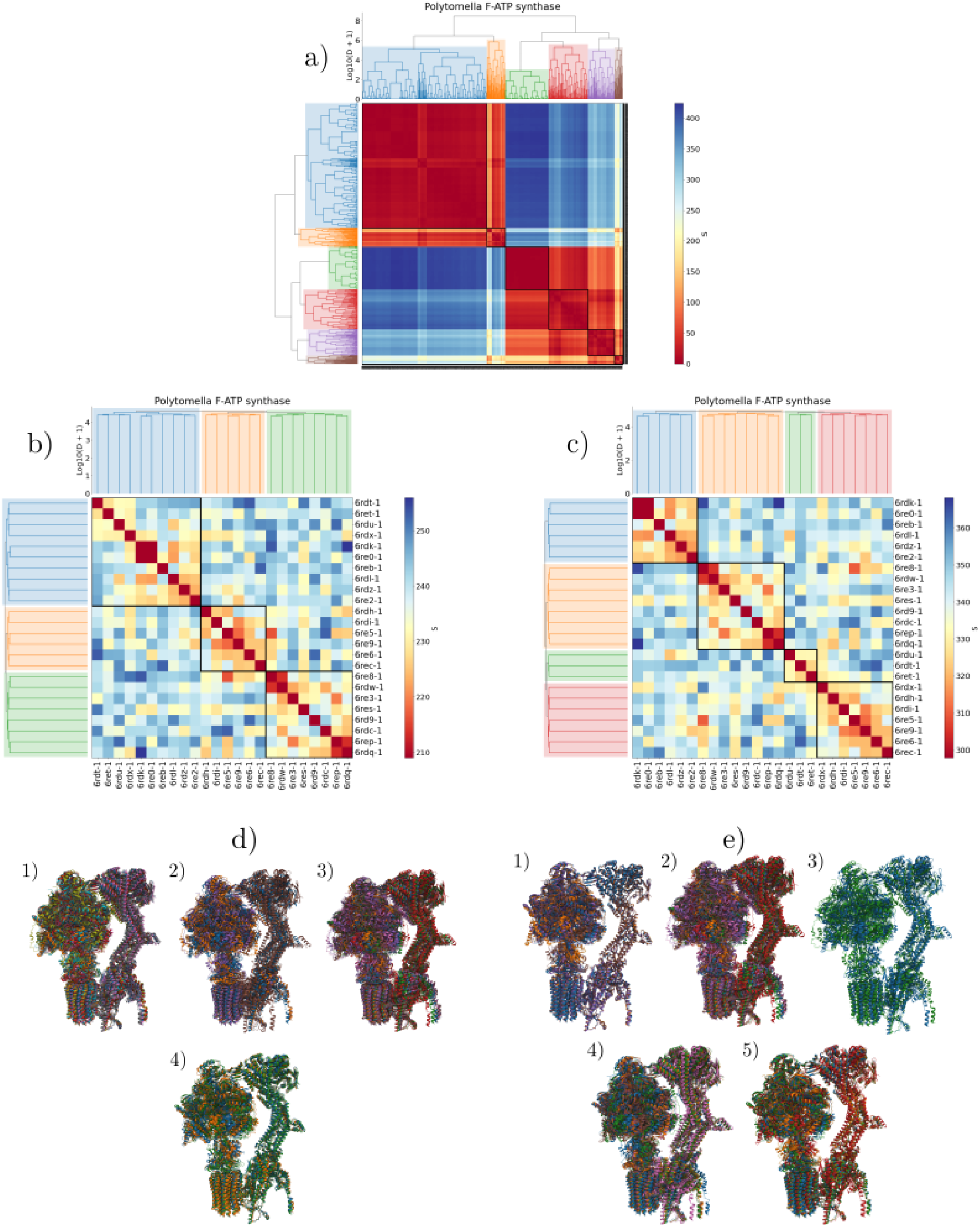
**a)** Chain-level conformational landscape, with the number of clusters determined using the second-order [24] difference statistic. **b)** Polytomella F-ATP score matrix and dendrogram, where clusters (1–3) are identified using the second-order difference statistic and arranged from top left to bottom right. **c)** Polytomella F-ATP distance-of-distances matrix and dendrogram, with clusters (1–4) determined using the standardized silhouette [8] score, also arranged from top left to bottom right. **d)** Superposed structures from clusters d1, d2, and d3 (top left to bottom right). d4 represents a superposition of one representative from each cluster. **e)** Superposed structures from clusters e1, e2, and d3 (top left to bottom right). e4 and e5 show a superposition of one representative from each cluster. Representatives and color bar calibration follow the same methodology as in Figure 2.

Classifying substates is even more challenging, yet they generally align with their respective cluster assignments despite variations in state definitions. This suggests that conformations exist along a continuum rather than as strictly distinct states. While the sparsity of the data makes it difficult to draw definitive conclusions, there is some agreement between the author annotations and FunCLAN’s clustering.

The superpositions in Fig. 3d further highlight that each cluster possesses distinct structural characteristics, particularly in the rotary domains and the “neck” connecting them. However, significant variation exists even within individual clusters. This is especially noticeable in the chains on the right side of the structures, which, from this perspective, fan in and out of the page. The complexity of conformational categorization is even more apparent when exploring the *.msvj files, which provide a clearer, more detailed visualization of these structural differences.

### Distances of Distances

Unlike the previous example, the “distances of distances” approach does not substantially reduce the number or severity of out-of-cluster hotspots. However, it does appear to make the clusters slightly more well-defined. In this case, we used standardized silhouette scores [8] to determine the optimal number of clusters, as the two different statistics yielded only two clusters.

In this case, we identify four clusters that contain fewer mixed conformations, suggesting that the clusters are more well-defined — at least according to author annotations. Similar to the previous example, closely related structures in one clustering approach remain closely related in the other. Notably, cluster 3 in the direct clustering and cluster 5 in the “distances of distances” clustering are identical.

There are several differences between direct clustering to “distances of distances” clustering:

● 6rdk, 6re0, 6reb, 6rdl, 6rdz, and 6re2 move from being part of cluster 1 to forming a new cluster 1.
● 6rdu, 6rdt, and 6ret shift from cluster 1 to form cluster 3.
● Cluster 2, along with 6rdx from cluster 1, merges to create cluster 5.

Although the two approaches yield different numbers of clusters, the underlying relationships between samples remain largely consistent. Notably, the “distances of distances” clusters appear more compact, suggesting that this method produces more well-defined groupings.

| Structure | Resolution | Conformation | Cluster |
| --- | --- | --- | --- |
| 6rdk | 3.7 | Rotary substate 1B | 1ll |
| 6re0 | 3.6 | Rotary substate 2A | 1lr |
| 6reb | 3.2 | Rotary substate 3A | 1rl |
| 6rdl | 3.7 | Rotary substate 1B | 1rrl |
| 6rdz* | 3.5 | Rotary substate 2A | 1rrrl |
| 6re2 | 3.2 | Rotary substate 2B | 1rrrr |
| 6re8 | 3.8 | Rotary substate 2D | 2lll |
| 6rdw | 3.8 | Rotary substate 1F | 2llr |
| 6re3* | 3.3 | Rotary substate 2B | 2lrl |
| 6res | 4.3 | Rotary substate 3C | 2lrr |
| 6rd9 | 3 | Primary rotary state 1 | 2rl |
| 6rdc | 3.2 | Primary rotary state 2 | 2rrl |
| 6rep | 3.1 | Primary rotary state 3 | 2rrrl |
| 6rdq | 4 | Rotary substate 1D | 2rrrr |
| 6rdu* | 3.5 | Rotary substate 1E | 3l |
| 6rdt | 3.4 | Rotary substate 1E | 3rl |
| 6ret | 4.3 | Rotary substate 3C | 3rr |
| 6rdx | 3.9 | Rotary substate 1F | 4ll |
| 6rdh | 3 | Rotary substate 1A | 4lrl |
| 6rdi | 3.2 | Rotary substate 1A | 4lrr |
| 6re5 | 3.2 | Rotary substate 2C | 4rll |
| 6re9 | 3.9 | Rotary substate 2D | 4rlr |
| 6re6 | 3.4 | Rotary substate 2C | 4rrl |
| 6rec* | 3.3 | Rotary substate 3A | 4rrr |

## Discussion

### What is a conformer?

This question lies at the heart of our work, yet a fully satisfactory answer remains elusive. One possible definition is that a conformer is a structural form that is biologically active in a way that is meaningfully distinct from other conformers. However, this leads to a deeper question: what qualifies as meaningfully different? The challenge in defining a conformer stems from the continuum of structural variations observed in biological systems. Some differences may be functionally significant, while others may represent minor fluctuations that do not alter biological activity. Determining the threshold at which a structural variation becomes a distinct conformer is a complex issue – one that this study seeks to explore.

The SARS-CoV-2 spike protein, for example, exhibits different biological activity depending on whether its receptor-binding domain (RBD) is in the “up” or “down” conformation. However, since the spike protein is a homotrimer, an interesting question arises: in the analyzed samples, why does the RBD transition occur in only a single chain, despite all three chains being structurally identical? These changes are not detected by FunCLAN because they do not affect the overall relationship between chains. From FunCLAN’s perspective, such variations do not constitute distinct conformations. This highlights a fundamental mismatch in the definition of “conformation” – while structural biology often considers RBD movement as a conformational change, FunCLAN only recognizes transformations that alter inter-chain relationships. As a result, what qualifies as a conformation in this context depends on external factors and the chosen analytical framework.

There is a way to make FunCLAN sensitive to these changes by segmenting the chains into separate regions, such as the RBD and the remainder of the chain. This would allow FunCLAN to detect these variations. However, this approach requires prior knowledge of the system being analyzed, making it less generalizable.

While existing methods like GESAMT can identify domains in single chains through pairwise comparisons, applying such an approach to protein complexes introduces significant computational challenges. The problem is inherently combinatorial, requiring an immense number of comparisons, and is highly susceptible to overfitting. Instead of yielding just the two expected domains, an algorithmic approach could generate an overwhelming number of small, fragmented domains. Without some yet-undiscovered insights, resolving this issue would demand an exhaustive search, likely leading to multiple equally valid partitions, all of which may still be overfitted.

We attempted this approach but ultimately recognized the sheer complexity of the task. Similarly, in the case of Polytomella F-ATP synthase, the three traditionally accepted conformations are best understood as discrete classifications of what is, in reality, a continuous conformational spectrum.

### Optimum number of clusters

The number of clusters is determined using data-driven heuristics, but heuristics nonetheless. While it is possible to specify a predefined number of clusters, this requires prior knowledge of the system, which is not always available. In this study, we consistently selected the method that yielded the highest number of clusters. Interestingly, in both direct clusterisations, this method was the second-order [24] difference statistic, whereas for the “distances-of-distances” clusterisations, it was the standard silhouette scores [8].

This choice, however, is inherently arbitrary, and it remains unclear whether one method is objectively superior to the other. The best approach must be evaluated on a case-by-case basis – much like the choice of a linkage function in hierarchical clustering. While different linkage functions exhibit distinct properties, and some are widely accepted as more suitable for specific problems, there is no universal best choice.

A fundamental challenge in clustering is its inherent sensitivity to small variations. Even minor fluctuations, such as those introduced by floating-point arithmetic, can yield different clustering results. When multiple samples receive identical scores, simply altering the order of input data can lead to different assignments. This sensitivity underscores the importance of avoiding overfitting when determining the number of clusters. Employing general, data-driven, and scale-agnostic heuristics is essential for drawing meaningful conclusions. This challenge is evident in Polytomella F-ATP synthase, where nearly identical samples are assigned to the same cluster but may not always be placed directly adjacent to one another.

### Limitations of considering chains as rigid bodies

Since chains are treated as rigid bodies, intra-chain changes may not be captured in the output if they do not affect the interactions between chains. This is intentional – FunCLAN was designed to identify global, complex-level conformations rather than small, localized changes. By prioritizing large, multi-chain transformations, the method inherently sacrifices sensitivity to minor, isolated variations within individual chains.

However, as we have seen, there are cases where the scientific community has agreed that certain small, local changes define distinct conformational states. In such instances, FunCLAN may fail to recognize them. Conversely, if such local variations are not functionally relevant to the overall conformation, FunCLAN remains robust, correctly categorizing them as insignificant. Whether FunCLAN aligns with conventional definitions of conformational change ultimately depends on the specific system being analyzed.

In cases where the assumption of chain rigidity is too strong and prior knowledge supports a different structural interpretation, users can manually preprocess structure files to redefine chains. For example, if a particular domain is known to undergo conformational changes while the rest of the chain remains largely fixed, the structure can be modified by splitting the chain into two separate entities – one for the static region and another for the dynamic domain. This allows FunCLAN to treat the domain as an independent chain, making it detectable as part of a larger conformational shift. Since such modifications require expertise in the specific protein being studied, they are best performed by researchers familiar with the system.

### Linkage functions

Agglomerative hierarchical clustering relies on a linkage function to determine how clusters are merged, with the goal of minimizing a specific criterion. The choice of linkage function significantly impacts the resulting clusters, as different functions are better suited for different types of data. FunCLAN employs the ALGLIB library and defaults to Ward’s linkage, as it prioritizes minimizing intra-cluster variation. However, users have the flexibility to select any linkage function supported by ALGLIB to better suit their specific analysis needs.

For those who wish to use alternative linkage functions or clustering algorithms, such as Direct Bubble Hierarchy Trees, FunCLAN provides an outputted score matrix in the form of a list of lists containing only the upper or lower triangular matrix above the diagonal. Users can apply their preferred clustering method to this matrix and then utilize the algorithm described in the Methods section to transform chain-level dendrograms into complex-level dendrograms. This allows for customized clustering while still maintaining compatibility with FunCLAN’s approach to complex-level conformational analysis.

### Future work

FunCLAN is more than just a C++ program – it is an integrative framework for analyzing protein complex conformations. To support this goal, the software suite includes a range of built-in analysis tools, allowing for comprehensive structural investigations. This paper serves as an initial demonstration of FunCLAN’s capabilities and the insights it can facilitate. For example, one of the sub-problems that FunCLAN solves is that of finding corresponding chains. This knowledge can be used to find chain interfaces and how they change between conformations.

## Supplementary Information

### PDBe Worker

There is currently a pipeline to execute FunCLAN on the PDBe database, but it is not currently in a regular schedule. Moving forward, it will focus on processing only updated complexes—those with modified, added, or removed assemblies. Initial test runs, conducted with 100 workers, have taken approximately 10 hours to complete. However, the primary computational burden stems from the substantial memory footprint required for full pairwise and multiple clustering. The pairwise comparison algorithm has a complexity of *O*(*N*^2^), even when avoiding redundant calculations. Hierarchical clustering requires *O*(*N*^2^) memory and *O*(*N*^2^*M*) computation time, where N represents the number of elements (samples) and M the number of features (dimensionality of the points). Consequently, only a few complexes with a large number of assemblies consume a disproportionate share of computational resources.

Furthermore, aside from matrix multiplication, the algorithms are strictly serial in nature. Fortunately, FunCLAN is written as two independent processes:

1. pairwise clustering for identifying chain parity between assemblies,
2. global clustering for conformational states.

It is feasible to develop two sub-programs that handle these tasks independently. This approach enables pairwise comparisons to run as distributed programs within a cluster. When a complex is updated — whether by modifying an existing assembly or adding/removing one — only the new entries need to be processed by the pairwise clustering algorithm. The global clustering can then integrate all pairwise results. This process must be carefully managed to prevent regressions, but it is entirely achievable. Ultimately, it allows for incremental updates to the analyzed data, making it a worthwhile endeavor.

## Methods

### Problem Definition

FunCLAN addresses the challenge of determining the number of distinct conformations within a set of macromolecular complexes that share a similar subunit composition and understanding how these conformational states differ. This concept is illustrated in Fig. 1, where differences between conformers are easily recognizable. However, in practice, this is a computationally complex problem. Each colored shape represents a distinct type of chain. The ultimate goal is to identify the different conformers and quantitatively assess the extent of their differences.

Levy et al. [6] demonstrated that complexes can be represented and classified using graph theory, an approach that has been applied to categorize protein complex families based on their graph structures. However, unless the connectivity between chains changes between conformers, the graph remains invariant. In Fig. 5, for example, a linear graph with six unique nodes represents a complex where each chain is a distinct node. One possible approach is to compare complexes as weighted graphs, but assigning appropriate weights requires prior knowledge of conformational changes, which varies across complex families. As a result, graph-based methods for recognizing conformational states rely on heuristics to account for these variations.

**Fig 5.**
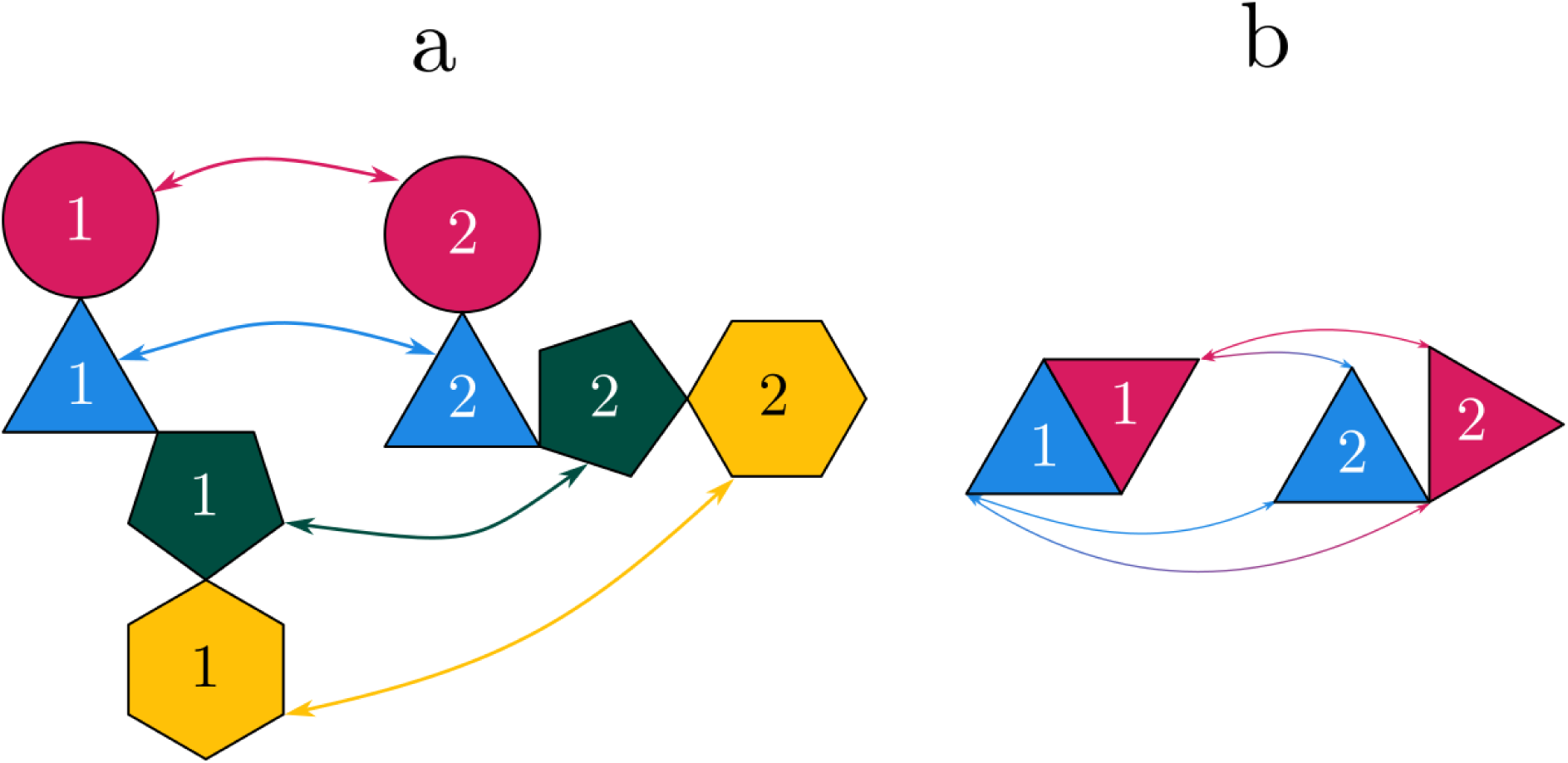
Matching chain pairs are not necessarily bijective. **a)** 1-to-1 match. **b)** *N* − *N* match.

Since topological solutions are either impractical or require prior empirical data, geometric methods are employed. While tools used for single-chain analysis [5] could be adapted, clustering complexes by conformational state introduces unique challenges that do not arise in single-chain analysis. For instance, every amino acid has an N- and C-terminus, allowing for a canonical direction to be defined for any given chain, which facilitates structural comparisons. However, this principle does not extend to chains within a complex, as there is no universally accepted method for ordering non-covalently linked chains. These chains can be parsed in any arbitrary order, adding complexity to the comparison process.

Similar to the single-chain problem, we must identify equivalent chains between complexes to enable meaningful comparisons. In the case of single chains, equivalences can be established at the level of individual amino acids [5] or secondary structures [4]. For complexes, these equivalents are entire chains. However, as noted in the previous paragraph, the lack of a standardized ordering for non-covalently linked chains presents a significant challenge in achieving this alignment.

Certain aspects of single-chain analysis can be disregarded when studying complexes. For instance, local conformational changes at the chain level, such as alterations in the shape of a binding site, are not the focus. Instead, the analysis concerns overall *structural* changes at the *complex level*, treating individual chains as rigid bodies.

The ultimate goal is to develop a robust, general, and automated method for classifying conformers of protein complexes while also quantifying their similarity or dissimilarity. Given these requirements, it became evident that a new approach was necessary.

### Corresponding chains

As previously stated, comparing samples requires having a map of chain correspondence between samples. FunCLAN does this based purely on sequence similarity. A different approach could use structural similarity, like q-score as defined in [5]. However, this adds extra computational cost without increasing accuracy and/or specificity because the complexes being analysed are the same.

For two chains to be paired, a threshold value, *t* ∈ [0, 1], of sequence similarity to the match sequence must be simultaneously met by both chains, Eq (1),

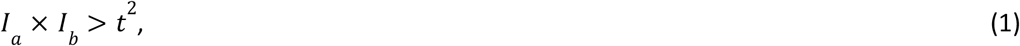

where *I*_*i*_ is the percentage of the sequence of chain *i*, found in the matched sequence, and *t* is the threshold value. This is equivalent to saying *I*_*a*_ > *t* and *I*_*b*_ > *t*, we do this rather than *I*_*a*_ > *t* or *I*_*b*_ > *t*, because we only want to pair up chains if both are sufficiently similar to one another. Otherwise, we may have a case where a small chain is a subset of a larger one. When this happens, the percentage similarity of the small chain to the matched sequence will be large, while the percentage similarity of the large chain to the matched sequence will be small. In fact, the small chain doesn’t even have to be a subset of the larger chain. The only requirement for this erroneous pairing to occur, is for the large chain to contain a sufficiently large percentage of the smaller chain’s sequence.

In the ideal example, where all chains are unique, as is the case in Fig. 5, the chain matching is bijective as in Fig. 6a, meaning each chain in either conformer has a single matching chain in the other.

In general, however, this is not always the case. A single chain in one conformer, may be matched to multiple chains in another, which happens any time a complex contains the same chain multiple times, for example, take the dimer in Fig. 6b. Such cases need to be disambiguated. For N-homomers, there can be up to *N*^2^ chain pairs. In order to solve the original problem, we need to define a *canonical* set of pairings. This is a challenging task, even for dimers. There is also the issue of missing or incomplete chains. In the following subsections – *Multiple matching chains* and *Incomplete or non-ideal data*–we address cases involving multiple chain pairs for a single chain and handling incomplete data, respectively. Regardless of the scenario, we begin by generating a list of potential chain pairs by comparing each chain in one sample to every chain in the other, retaining only those that meet the threshold criteria. This list serves as the foundation for the rest of the algorithm. For now, we assume an ideal case where each chain pairs uniquely, all chains are represented, and every pair meets the required similarity threshold across all samples.

### Chain-level clustering

To illustrate our algorithm, we use a simple example shown in Fig. 1. Here, we have three samples of a single complex, composed of unique chains represented by distinct shapes and colors. Each chain is uniquely identified by its shape, color, and inscribed number.

Since the chains are all distinct, we define a bijective map from any given chain in one sample, to its equivalent in a different one. We then define a transformation that best superposes matching chain pairs to one another. We call this transformation from chain *j* to *i* as *T*_*i*,*j*_.

If we go back to Fig. 1, we can see a few things. If we compare samples 1 and 2, we can identify two sets of chains whose internal relationships are conserved. These being the sets of {○, △} and {⬠, ⬡}. It’s the relationship between these two sets that changes. Similarly, if we compare samples 2 and 3, we can identify that the yellow hexagons are in roughly the same position, just shifted slightly between samples.

The key insight is that our goal is not to compare chains, but to compare transformations. We aim to assess how similar these transformations are to one another. By identifying equivalent transformations or sets of transformations, we can detect concerted movements between conformers. Measuring the similarity of these movements allows us to understand how the relationships between chains evolve or remain conserved as the system transitions from one configuration to another.

We can extend this idea to define a score that quantifies the similarity between two transformations. However, this score must satisfy several key properties. Let’s define two superposition transformations, *A* and *B*, which include both the rotation matrix and translation vector.

1. Comparing transformation *A* to *B* must be the same as comparing *B* to *A*, therefore *s_A,B_ = s_B,A_*
2. Self-comparison must have a score of zero, **s*_A,A_* = 0.
3. The bigger the difference, the larger the score.
4. Comparing equivalent transformations (i.e. their inverses) must yield the same result, 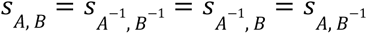. This is because *A*: *c*_2_ ⇒ *c*_1_, which is to say any transformation *A* superposes chain *c* of sample 2 onto chain *c* of sample 1, should be equivalent to *A*^-1^: *c*_1_ ⇒ *c*_2_ which superposes chain *c* of sample 1 onto chain *c* of sample 2.

This property ensures the score is invariant to swapping fixed and moveable chains.

Our requirements mean we need a distance matrix. Our only special provision is number 4. Indeed, the score can be defined quite elegantly. Let 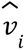 be the canonical basis vector for dimension *i* in 3D Euclidean space. We can define inverse transformations *C* = *A*^-1^ and *D* = *B*^-1^ and treat transformations as functions that act on our basis vectors, which lets us define our score as follows:

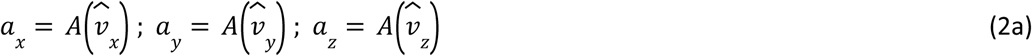

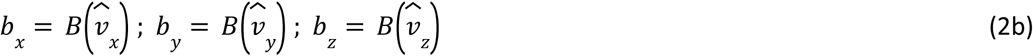

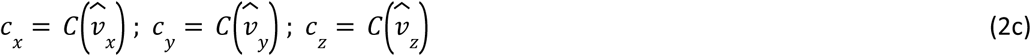

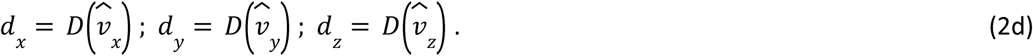

The vectors *a_i_*, *b_i_*, *c_i_*, *d_i_* are the transformed unit vectors. These are by themselves not very useful, but we can compute an average displacement (RMSD) between a transformed vector and its corresponding basis vector, and place them into a characteristic vector for each transformation.

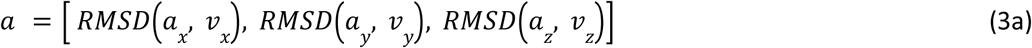

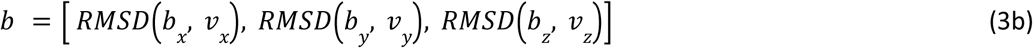

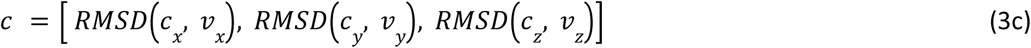

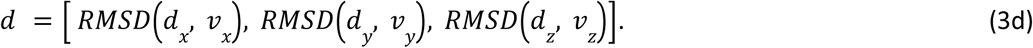

These new vectors give us the average displacement each transformation imposes on each basis vector. We can then get the average difference (RMSD) between the vectors corresponding to *A* and *B*, as well as their inverses as per condition number 4 of our scoring function. We have four of these, *A* to *B*, *A* to *B*^-1^, *A*^-1^ to *B*, and *A*^-1^ to *B*^-1^, hence our score is the comparison which yields the smallest value.

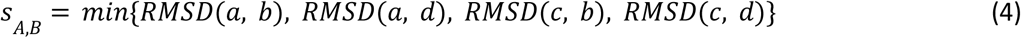

This approach allows us to measure the difference in the *effects* of the transformations, rather than the transformations themselves. It inherently accounts for both rotational and translational components, and possesses all the desired properties. While other distance measures could be used, the RMSD is a natural choice in this context. Additionally, although it is possible to modify the transformations to exclude either the rotational or translational component, we do not see a relevant use case for this in our application.

Now, we take our list of paired chains and determine the transformations that superpose each pair. We then compare each transformation to all others using our scoring function (Eq. 4). Since the distance matrix is symmetric and has a zero diagonal, we only need to compute the upper or lower triangle, excluding the diagonal. This reduces the number of operations from *N*^2^ to *N*(*N* − 1)/2, where *N* is the length of the paired chain list.

With the score matrix in hand, we now need a method to identify and measure relationships within the data. Since our score matrix is a distance matrix, we can directly apply it in hierarchical agglomerative clustering to analyze the data. Applying this approach to our example in Fig. 1a would generate dendrograms similar to those in Fig. 1b-1d.

Fig. 1b-1d shows pairwise comparisons at the chain level, but our goal is to compare *all* complexes at the *complex* level.

Once we have our pairwise comparisons, we can clusterise all pairwise transformations, producing a result like Fig 1e, which is based on a distance matrix reflecting the conformational landscape at the chain level. However, our objective is to extract complex-level information..

### Whole complex clustering

To clusterise based on complexes we need a way to summarise or recover their contributions to the conformational landscape. Luckily, we already have all the necessary information in the distance matrix, which compares all individual chains.

If we index the matrix by the dendrogram, the dendrogram leaves correspond to columns/rows of the matrix. We can then identify all the leaves containing a transformation that uses a chain from sample *i*. Since the matrix is indexed by the dendrogram, these leaves correspond to the relevant columns and rows in the distance matrix. Next, we concatenate all the columns and rows into a single vector, *p*_*i*_, which encodes the pairwise relationship of sample *i* to all other samples. By doing this for all samples, we can compare these vectors to each other. The function we use for this is again the RMSD,

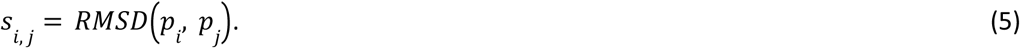

This approach allows us to create high dimensional points, or *conformational coordinates*, for each sample. We use these scores to perform another round of hierarchical clustering, this time based on pairwise comparisons at the whole complex level. Since the distance matrix is indexed by the dendrogram and is symmetric, the ordering of the dendrogram’s ordering can change without affecting the score. This is important when the input order results in two leaves to switch places.

Each row *S_i_* (or column) represents sample *i*’s relationship to every other sample, forming matrix *S*, which captures only inter-sample relationships. However, if we view each row/column as a multidimensional point representing a sample’s relationship to all others, this perspective aligns with an idea from portfolio optimization [7], where it helps mitigate clustering overfitting and enhances robustness. By recognizing that each row/column is itself a multidimensional point that represents a sample’s relationship to every other sample, we can apply a similar transformation as in the previous step. Specifically, we redefine 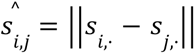, using Euclidean distance rather than RMSD. The choice of Euclidean distance prevents artifacting without inflating the score as more samples are added, this is because the scores of the previous clusterisation have already been normalised. This “distances of distances” matrix, 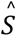, effectively captures how each sample perceives the overall conformational space. This measure is more robust and global, as it reduces out-of-cluster hotspots—regions of high similarity that lie outside defined clusters. These hotspots arise when pairwise similarities between samples from different clusters are stronger than their similarities to other members within their own clusters. This issue is inherent to hierarchical clustering, which optimizes an ensemble-wide objective function rather than individual pairwise distances. High-dimensional, sparse, and/or non-linear problem spaces are particularly susceptible to such discrepancies, where two structures may be highly similar in some respects while also closely resembling an ensemble average of many others. In practice, both approaches yield similar and reasonable results, but the distances-of-distances method reduces both the number and intensity of out-of-cluster hotspots, leading to a more stable clustering outcome.

A key issue with this approach is that two points with high scores may be clustered together even if they are fundamentally different, simply because their transformations are distinct from all others, including each other. One way to address this is by penalizing scores when values are high or exhibit large dispersion. However, this introduces heuristics that may not be universally applicable. This challenge reflects the high dimensionality and sparsity of the problem. Essentially, we are projecting samples from an extremely high-dimensional conformational space into a much lower-dimensional representation. As a result, distinct conformations may appear similar because their relative distances to other points in the dataset are comparable. This limitation stems from an insufficient amount of data to fully capture the conformational landscape, and we must acknowledge it as an inherent trade-off.

This approach requires all samples to have the same number of equivalent chains, as comparisons are only valid when points share the same dimensionality. If a sample *i* is missing a chain, it will have fewer comparisons than other samples, resulting in its point *p*_*i*_ having *N* missing entries in the *N* × *N* matrix. Similarly, if a chain is incomplete or falls below the similarity threshold, the point will lack the same number of entries as the others. Since hierarchical clustering relies on a global measure, a single non-conforming sample can cause the procedure to fail. To prevent this, FunCLAN detects such cases early and provides a detailed error message indicating the number of valid comparisons per sample. The process must then be rerun without the problematic samples. Alternatively, non-strict mode, explained in the *Incomplete or Non-Ideal Data* section, can be used to handle such cases.

### Multiple matching chains

As mentioned in *Corresponding chains*, we often pair the same chain with multiple other chains. Resolving this issue is relatively straightforward, but it must be addressed before proceeding with the multiple comparison. It is important to keep a few key points in mind.

● Each transformation involves two chains, a fixed and a moveable one.
● The dendrogram is ordered by the ascending height of each cluster.
● Each height is associated with a cluster.
● Each cluster can have:

○ Two transformations, they are more similar to each other than anything else.
○ One transformation, the transformation is similar to a set of transformations.
○ No transformations, sets of transformations are similar to each other.
● All fixed chains belong to the same complex, all moveable chains belong to the other complex.

The algorithm is as follows.

1. Initialise an empty list of valid transformations, and a list of past chain pairs.
2. Loop over the dendrogram starting from the smallest height to the largest.
3. If the cluster is made up of two transformations.

a. Check whether the *fixed* or *moveable* chain used by the transformation on the *left* side of the cluster is in the list of past chain pairs.

i. If neither of them are in the list, the *left* side of the cluster is flagged as potentially valid.
b. Check whether the *fixed* or *moveable* chain used by the transformation on the *right* side of the cluster is in the list of past chain pairs.

i. If neither of them are in the list, the *right* side of the cluster is flagged as potentially valid.
c. If the *left* side is potentially valid; and the right side is potentially valid; and the *fixed* chain of the transformation in the *left* side is *different* to the *fixed* chain of the transformation in the *right* side; and the *moveable* chain of the transformation in the *left* side is *different* to the *moveable* chain of the transformation in the *right* side.

i. Add both transformations to the list of valid transformations.
ii. Add both pairs of chains to the list of past chain pairs.
d. If the *left* side is potentially valid; and the *right* side is *not* potentially valid.

i. Add the transformation on the *left* side of the cluster to the list of valid transformations.
ii. Add the pair of chains corresponding to the transformation on the *left* side of the cluster to the list of past chain pairs.
e. If the *left* side is *not* potentially valid; and the *right* side is potentially valid.

i. Add the transformation on the *right* side of the cluster to the list of valid transformations.
ii. Add the pair of chains corresponding to the transformation on the *right* side of the cluster to the list of past chain pairs.
4. If a cluster is made up of one transformation and another cluster. Only check the side of the cluster which points to a transformation.

a. Check whether the *fixed* or *moveable* chain used by the transformation on this side of the cluster is in the list of past chain pairs.

i. If neither chain is in the list.

1. Add the transformation to the list of valid transformations.
2. Add the pair of chains corresponding to the transformation to the list of past chain pairs.
5. If a cluster contains no transformations, move on to the next height.

By the end of this process, we obtain a subset of the most similar transformations, ensuring each chain appears only once. We then cluster them using our scoring function to generate the definitive dendrogram for pairwise comparison. Multiple clustering considers only the valid transformations from each pairwise comparison.

### Incomplete or non-ideal data

As mentioned earlier, points must share the same dimensionality for comparison. We can enforce this requirement by standardizing all points to the same dimensionality. If the matrix comparing all complex chains has a dimensionality of *N* × *N*, we define each sample’s points to have a dimensionality of *N* × 1. Instead of concatenating columns into a single point, we elementwise add them into the corresponding *N* × 1 point.

While this reduces resolution, it ensures uniform dimensionality across all points, regardless of data quality.

### Whole complex-complex superposition

FunCLAN derives a whole-complex superposition transformation for pairwise comparisons by preserving the alignments of all valid superpositions. It concatenates these alignments into a single synthetic superchain, which is then used to superpose both complexes in their entirety, treating individual chain alignments as the matched sequences. This approach eliminates the need for realignment and prevents artefacts that could arise from *de novo* alignment of superchains. Additionally, it ensures invariance to the order of chain concatenation — an issue that can arise when computing new alignments on superchains.

### Determining the number of clusters

We have two data-driven heuristic methods:

1. second-order difference statistic [24],
2. standardised silhouette score [8],

for determining the optimum number of clusters. For the purposes of this paper, we assume the clusters represent conformers, but this may not always be true. This is especially relevant in two cases:

1. When there is chaining/laddering in the dendrogram, since that means there is very little difference between clusters.
2. When the *inter*-cluster differences are similar to the *intra*-cluster differences.

This is why this part is performed by an external post-processing Python script rather than FunCLAN itself. Information on the methods can be found in their respective publications.

### Method comparison

Both methods, described above, are scalable and robust. Whether the score matrix or dendrogram operates on vastly different scales, they provide informed best-fits for the number of distinct clusters represented in the data, based on their respective objective functions. However, both are data-driven heuristics, completely agnostic to the underlying meaning of the dendrogram. Since they rely on clustering results, they are sensitive to the same factors that influence the clustering method to which they are applied.

Both methods have different objective functions, which means that given the same distance matrix and clustering results, both methods may yield a different number of optimal clusters. Generally, the second-order [24] difference statistic splits the dendrogram into more clusters. This is the default method in our post processing script.

More clusters do not necessarily lead to a better solution, so it’s important to test both approaches while remembering that they are data-driven heuristics, not definitive answers. Users can also specify the number of clusters manually, but this should only be done by those with in-depth knowledge of a given complex.

In the samples presented in this paper, there is little to no difference when analyzing clustering at the complex level. Beyond these examples, we have applied FunCLAN to a dataset of over 350 distinct complexes, and this trend largely holds. The most significant differences in the optimal number of clusters arise when both methods are applied to chain-level dendrograms. However, this falls outside the scope of this paper. That said, chain-level information may be valuable to other researchers. It has already proven useful for studying interfaces, where identifying corresponding chain pairs is essential for constructing both chain-level and complex-level dendrograms. This research is currently ongoing.

For this reason, we believe that providing two different methods for finding the optimal number of clusters is a good idea. It is also possible for researchers to create custom methods by adapting these ideas in the algorithms we have presented.

### Picking Cluster Representatives

It may be useful to have a reliable method for selecting cluster representatives. Here we propose a simple, doubly-centered, and robust measure.

If there are *M* structures in a cluster, then there are *M* vectors (one for each cluster member) of length *M* − 1 (removing the self-comparison because it is zero), a sample’s score is the median of its vector. The entire cluster has (*M* − 1)/2 unique, non-zero scores (the cluster sub-matrix is symmetric with zero diagonal), the cluster’s score is the median of these scores. We then compare the cluster score to the score of each of its *N* members. The member whose Euclidean distance is closest to the overall cluster median is taken as the representative. While this is just one of many possible methods for choosing a representative, it is a simple, doubly-centered, and robust measure, making it particularly appealing. Alternatives to the median could include the mean, standard deviation, or other statistical measures.

### Generalising FunCLAN

Generalizing FunCLAN is straightforward. For instance, if a protein consists of a single chain with multiple secondary structures, a GEMMI-supported file, where each secondary structure is treated as a separate chain, can be created. Applying this to all samples enables FunCLAN to analyze these “chains” individually. The same approach can be extended to domains or even custom-defined regions.

